# Habitat percolation transition undermines sustainability in social-ecological agricultural systems

**DOI:** 10.1101/2021.06.07.447347

**Authors:** Diego Bengochea Paz, Kirsten Henderson, Michel Loreau

## Abstract

Steady increases in human population size and resource consumption are driving rampant agricultural expansion and intensification. Habitat loss caused by agriculture puts the integrity of ecosystems at risk, and threatens the persistence of human societies that rely on ecosystem services. We develop a spatially explicit model describing the coupled dynamics of an agricultural landscape and human population size to study the effect of different land-use management strategies, defined by agricultural clustering and intensification, on the sustainability of the social-ecological system. We show how agricultural expansion can cause natural habitat to undergo a percolation transition leading to abrupt habitat fragmentation that feedbacks on human’s decision making, aggravating landscape degradation. We found that agricultural intensification to spare land from conversion is a successful strategy only in highly natural landscapes, and that clustering agricultural land is the most effective measure to preserve large connected natural fragments, avoid severe fragmentation, and thus, enhance sustainability.

## 1 Introduction

In recent decades, the global human population has continued to grow, and so has the mean per capita food consumption (Barrett *et al*., 2020). The increased demand for resources has led to worsening environmental degradation, such that food security and nature preservation are major concerns for human societies worldwide. Both zero hunger and environmental conservation are among the United Nations’ Sustainable Development Goals (UN, 2015). Furthermore, food security and nature are interdependent, as nature provides essential ecosystem services to agricultural systems (Power, 2010). Even though it has been pointed out that ending hunger is a matter of resource accessibility more than resource availability (Chappell & LaValle, 2011; Tscharntke *et al*., 2012), it is widely assumed that increasing agricultural production is the key to achieving food security in the future (Chappell & LaValle, 2011). However, agriculture is a major driver of environmental degradation, therefore increasing agricultural production will undoubtedly come with an environmental cost (Kehoe *et al*., 2017). This raises questions about the trade-offs between the zero hunger and environmental conservation goals, as they may counteract each other. Achieving global food security while preserving the environment is thus one of this century’s key sustainability challenges.

Increases in food production will require either agricultural expansion or intensification. Agricultural expansion relies on the cultivation of larger areas of land to achieve greater food production, while agricultural intensification generally relies on greater synthetic inputs to increase production per unit area. Agricultural expansion is today’s main threat to biodiversity conservation as it directly causes habitat loss and fragmentation (Tilman, 1999), but agricultural systems can also contribute to the conservation of biodiversity (Dudley & Alexander, 2017). Low-intensity, wildlife-friendly farming (Perfecto & Vandermeer, 2010; Chappell & LaValle, 2011) is one such method for combining food production and conservation. However, low-intensity agriculture requires larger land surfaces to achieve the same production. The expansion of agricultural land could encroach on ecosystems that serve as habitats for wild species (Bengtsson *et al*., 2005; Perfecto & Vandermeer, 2010). Furthermore, it has been shown that the biodiversity in wildlife-friendly agricultural land is smaller than in natural land, making it unclear whether low-intensity practices can compensate for changes in land use (Phalan *et al*., 2011; Balmford *et al*., 2019). Rather than mixing agriculture with biodiversity conservation, agricultural intensification has been suggested as a sustainable solution to increase food production while sparing natural land from conversion. However, conventional intensification relies on heavy use of fertilizers, pesticides, artificial irrigation and machinery, all of which foster the degradation of the cultivated land as well as nearby habitats and freshwater systems by spillover (Tscharntke *et al*., 2012).

Without integral land use management policies at national and international scales or a clear consensus on the land sparing-sharing debate, the choice of expansion or intensification is based on food production and economic gains (Lambin & Meyfroidt, 2011; le Polain de Waroux *et al*., 2016). At present, the majority of land suitable for agricultural expansion is located in tropical regions (Byerlee *et al*., 2014). Thus tropical forests and grasslands in Africa, Asia and Latin America currently face rampant deforestation, putting at risk the integrity of some of the world’s species-richest ecosystems. On all three continents, agricultural expansion and intensification are occurring simultaneously in response to the soaring global food demand and fluctuations in international markets. The disastrous environmental consequences of such dramatic changes in land use are already present and are expected to worsen if nothing changes (Tölle *et al*., 2017; Ordway *et al*., 2017; Boers *et al*., 2017; Stoy, 2018; Staal *et al*., 2020; Ruiz-Vásquez *et al*., 2020; Baldassini & Paruelo, 2020).

Taubert *et al*. (2018) assessed the current state of forest cover across the Asian, African and Latin American tropics through an analysis of satellite images, and concluded that tropical forests may be close to a percolation transition (Aharony & Stauffer, 2003). A percolation transition occurs when progressive habitat loss causes an abrupt increase in landscape fragmentation when the amount of habitat drops below a certain threshold. In practice, this leads to the disappearance of large-sized connected habitat fragments, which are replaced by many smaller ones.

Habitat fragmentation may initially increase the flow of ecosystem services to human-transformed systems by expanding their edges with natural land fragments (Mitchell *et al*., 2015b). However, as habitat patches become smaller, more isolated and with a larger edge-to-area ratio, the deleterious effects of habitat fragmentation on ecosystem functioning grow, to the point where fragments become too small to provide ecosystem services to the surrounding area (Haddad *et al*., 2017). Long-term experiments have shown that fragmentation leads to the degradation of crucial ecosystem services for agricultural production such as nutrient retention and pollination, increases the vulnerability of natural systems and threatens their persistence (Haddad *et al*., 2017). When faced with a decline in agricultural production associated with a decrease in regulating and supporting services land managers are likely to turn to crop-land expansion to compensate their losses, causing further habitat loss. Alternatively they may attempt to increase yields via intensification, furthering the degradation of habitat quality. Current agricultural practices raise many sustainability concerns, especially when habitat loss can generate a sudden percolation transition, where a large connected fragment is divided into many small fragments by removing a few key connections.

Achieving food security while preserving the environment requires informed and careful land use policies and management. The large spatial and time scales at which landscape and social processes occur make it difficult to identify paths towards sustainability using empirical studies alone. Theoretical and modeling approaches provide useful perspectives that shed light on the possible outcomes of different future scenarios for land use management and policy. Coupled human-nature dynamical models are of particular interest to tackle these questions as they explicitly take into account the bi-directional feedbacks between the environment and human societies (Motesharrei *et al*., 2014; Balbi *et al*., 2020). Models that account for population dynamics (Lafuite *et al*., 2017, 2018; Cazalis *et al*., 2018; Henderson & Loreau, 2019, 2020; Bengochea Paz *et al*., 2020) are particularly suitable to study long-term dynamics, as changes in population size greatly affect societal pressures on the environment. However, within the current body of literature there are no models accounting for human demography and land dynamics that explicitly account for spatial structure.

The aim of the present work is to understand the implications of uninformed land-use management on the sustainability of social-ecological agricultural systems. We build a model coupling human demography with spatially explicit land-cover dynamics to study the influence of different management practices in landscape structure and resource production. We then use percolation theory to provide a mechanistic understanding of the causes behind abrupt landscape fragmentation and analyze the effects on ecosystem services provision and agricultural production. Additionally, we illustrate how bidirectional feedbacks between natural and human systems can trigger naive decision-making in the wake of a habitat percolation transition that traps the system in a path of social-ecological collapse. Finally, we focus on agricultural intensification and spatial planning as potential measures to enhance the sustainability of agricultural landscapes. Furthermore, we study the likelihood of their success depending on the initial landscape composition. Our work sheds light on how to better design agricultural land-use management policies for conservation and sustainability purposes, as consumption demands and the human population continue to grow.

## 2 Model and Methods

### Model overview

We formulate a spatially explicit stochastic model describing the coupled changes of human population density and the landscape humans exploit to produce resources. We model the landscape as a lattice with periodic borders where the state of each cell is determined by its land use/cover type. We consider four types of land use/cover: natural, degraded, low-intensity agriculture and high-intensity agriculture. Transitions between land use/cover types are stochastic and their probabilities per unit time, hereafter called propensities, are determined either by ecosystem services provision, which is a function of the landscape’s state, or by decision-making. In turn, human population dynamics and decision-making depend on the resources per unit time yielded by the landscape, which are also dependent on the ecosystem services provided to agricultural cells.

In what follows, all the model equations and mathematical expressions for the transition propensities are presented in their non-dimensional form. The details for the full derivation of the equations and the non-dimensionalization can be found in Appendix 1.

### Measure of ecosystem services provision

Ecosystem services are quantified by the size of connected natural fragments and they flow to neighboring cells. Following (Mitchell *et al*., 2015a) we assume that the magnitude of ecosystem services generated by a fragment saturates with its area. We opt for a power-law function with an exponent smaller than one to describe the saturation. Assuming that the flow of ecosystem services is limited to the closest neighbors of a natural cell yields the following expression for the total amount of services *ε_i_* in a given cell *i*

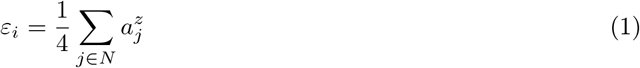

where the sum takes into account all the natural cells *N* of the landscape, and *α_i_* is the relative area of the natural fragment to which cell *j* belongs. *z* < 1 is the saturation exponent. We apply a Von Neumann neighborhood to constrain the non-dimensional ecosystem services provision within the interval [0, 1] using the 1/4 normalization factor (see Appendix 1 for details).

### Resource production

The total amount of resources perceived by the human population is the sum of the production per unit time of all agricultural cells in the landscape. In order to account for the contribution of ecosystem services to agriculture, we assume that the productivity of low-intensity agricultural cells is a linear function of the ecosystem services they receive. On the contrary, we assume the productivity of high-intensity agricultural cells is constant and independent of ecosystem services provision. The equation for total resource production per unit time *Y* is

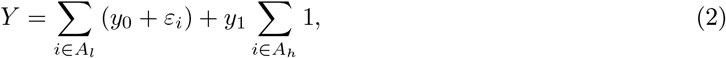

where the sums are over the sets of low-intensity *A_l_* and high-intensity *A_h_* agricultural cells and *y*_0_*y* and *y*_1_ are non-dimensional parameters representing the baseline productivity of low-intensity agriculture and of high-intensity agriculture (see Appendix 1 for details).

### Human demography

We assume that human population density *P* follows deterministic logistic dynamics with a carrying capacity that evolves over time subject to changes in resource production. We define the human carrying capacity as the maximum population density that can be supported for a given resource production *Y* and per capita consumption per unit time. Assuming a constant per capita consumption per unit time yields the following non-dimensional equation for the population density:

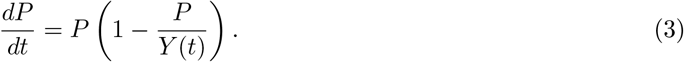

Note that the per capita consumption per unit time does not appear explicitly in the equation as it is encapsulated in the non-dimensionalization of the population density (see Appendix for details).

### Agricultural land use management

We consider two land use transitions related to agriculture: expansion and intensification. Expansion is defined as the transition from a natural state to low-intensity agriculture and intensification is the transition from low-intensity to high-intensity agriculture. We assume that the expansion *π_E_*(*i*) and intensification *π_I_*(*i*) propensities at cell *i* grow linearly with the human population’s perceived lack of resources Δ*r* = *p* − *Y*. The equations for expansion and intensification, assuming that both practices occur with uniform probability in space, are:

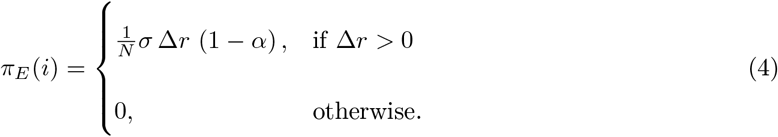

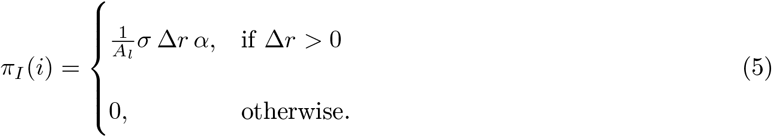

*N* and *A_l_* are the total number of natural and low-intensity agricultural cells in the landscape respectively. Parameter *α* ∈ [0 : 1] controls the preference for agricultural intensification and *σ* represents the population’s sensitivity to resource deficiency.

To study the influence of the spatial configuration of agriculture in the landscape we introduced a clustering parameter *ω* that controls the likelihood of expanding or intensifying agriculture next to cells with the same type of land cover (see Appendix 1 for details on the formalization).

### Loss of agricultural land

We consider fertility loss due to soil erosion as the main driving factor of agricultural land degradation (Pimentel, 2006). Urban expansion over fertile agricultural land is also an important cause of current cropland loss, however we do not account for this mechanism since we do not model human settlements in a spatially explicit way. We assume that the average time to fertility loss is a function of the amount of ecosystem services an agricultural cell receives. A large amount of ecosystem services contributes to maintaining fertility over longer periods of time. The propensity *π_L_*(*i*) of a fertility loss transition in agricultural cell *i* is given by:

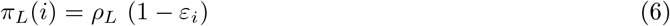

where *ρ_L_* is the sensitivity of fertility loss to ecosystem services provision and represents the decrease in rate per unit of ecosystem service of the fertility loss propensity. In addition, we assume that low-intensity agricultural cells transition back to a natural state when not in use; whereas high-intensity cells transition to a degraded state, as a result of the fertility loss transition, given that intensification has a greater impact on the soil.

### Passive land recovery and degradation

We call land recovery (degradation) the transition from a degraded (natural) to a natural (degraded) state without human intervention. We assume that the propensity of a recovery (degradation) transition grows (diminishes) with ecosystem services provision (Cramer *et al*., 2008). The land recovery *π_R_*(*i*) and degradation *π_D_*(*i*) propensities in cell *i* are:

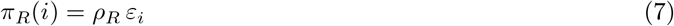

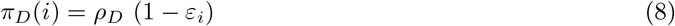

where *ρ_R_* and *ρ_D_* are land recovery and land degradation sensitivities to ecosystem services provision, respectively. Passive recovery and degradation transitions allow for the propagation or the containment of human induced perturbations on the landscape.

### Computational implementation and numerical simulations

We simulate the social-ecological dynamics in continuous time using the Gillespie’s Stochastic Simulation Algorithm (Gillespie, 1977) coupled with a Runge-Kutta 4 solver for the population density differential equation (see Appendix for details). The size of the landscape lattice is 40×40 cells (1600 pixels) and border conditions are periodic. Model runs were replicated to account for the inherent stochasticity in the model. All the code is open access and released in the repository https://doi.org/10.5281/zenodo.4905944.

We focus our study on the impact of land-use management practices on the sustainability of the social-ecological system. More precisely, we address the effect of the sensitivity to resource deficiency, agricultural intensification and agricultural clustering on landscape structure and study how landscape fragmentation can undermine the social-ecological system’s sustainability.

**Table 1:** Description of the model’s non-dimensional parameters and numerical values used for the simulations presented on this article.

| Parameter | Interpretation of the non-dimensional parameters | Values |
| --- | --- | --- |
| $z$ | Saturation exponent of ecosystem services provision with the area of a natural fragment | 0.25 |
| $y_1$ | Yield of intense agriculture relative to the yield contribution of ecosystem services to low-intensity agriculture | 1.2 |
| $y_0$ | Baseline yield of low-intensity wildlife friendly farming relative to the yield contribution of ecosystem services to low-intensity agriculture | 0.2 |
| $\alpha$ | Preference for agricultural intensification over agricultural expansion | [0.0 – 1.0] |
| $\omega$ | Clustering of agricultural land | [0.0 – 10.0] |
| $\sigma$ | population's sensitivity to resource deficiency: increase rate of the expansion and intensification propensities per unit of resource deficiency | [1 – 100] |
| $\rho_L$ | Fertility loss's sensitivity to ecosystem services provision: decrease rate of the fertility loss propensity per unit of ecosystem services | 0.02 |
| $\rho_R$ | Land recovery sensitivity to ecosystem services provision: increase rate of the land recovery propensity per unit of ecosystem services | 0.2 |
| $\rho_D$ | Land degradation sensitivity to ecosystem services provision: decrease rate of the land degradation propensity per unit of ecosystem services | 0.02 |

## 3 Results

### Agricultural intensification: a double-edged sword

We study the temporal changes on the social-ecological system starting from low population densities at equilibrium with agricultural production in an almost pristine landscape. Departing from the nearly pristine nature initial conditions, spontaneous land cover fluctuations drive the dynamics as they constantly push the system away from the equilibrium. Land cover changes can diminish ecosystem services provision and resource production leading to either adjustments in population density or to agricultural expansion. In the absence of agricultural intensification, the system results in collapse-recovery cycles or a sustainable steady state depending on the population’s sensitivity to resource deficiency (Figure 1a & b).

**Figure 1:**
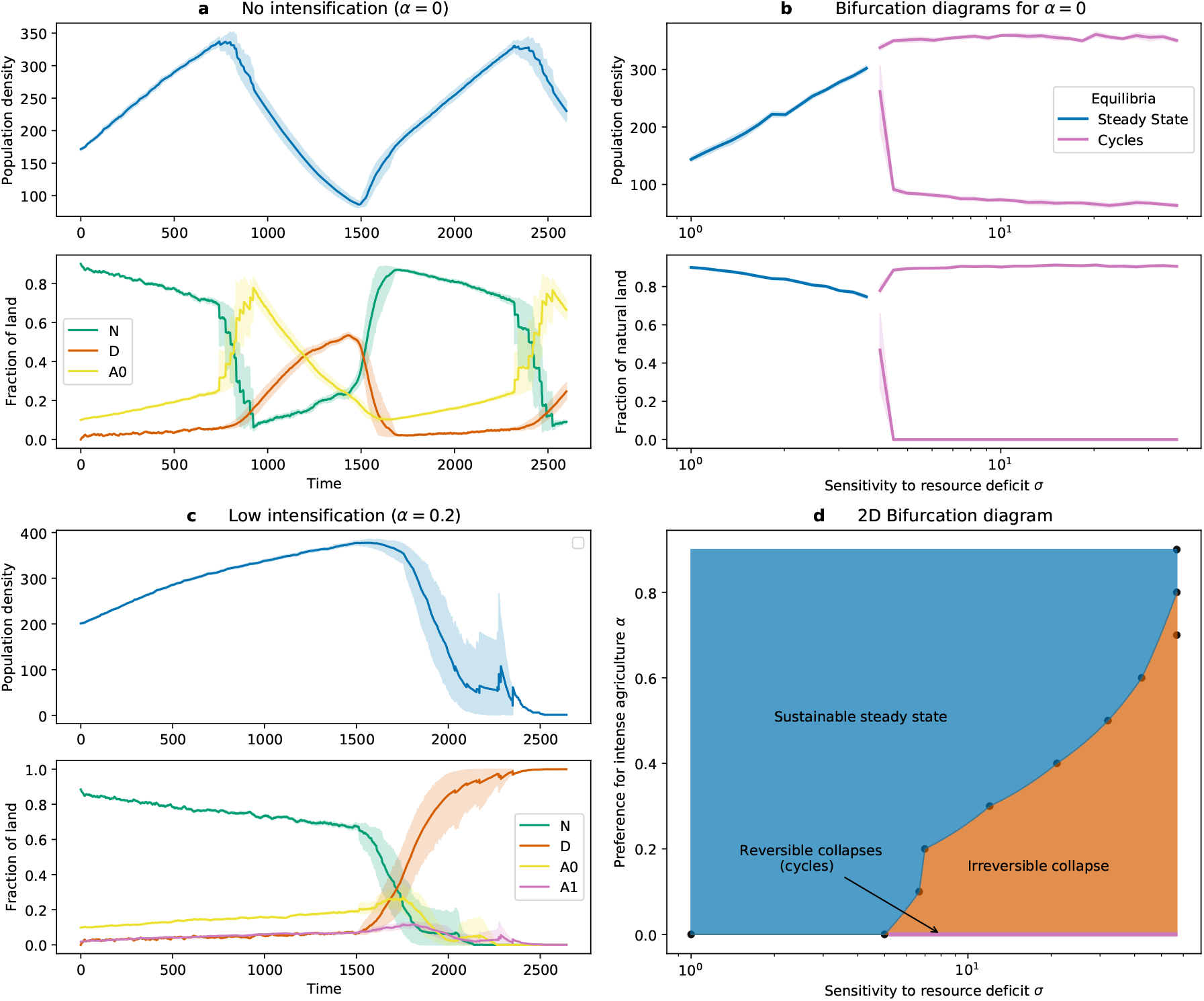
Effect of agricultural intensification on the temporal dynamics and the long-term states of the system. a) Collapse-recovery cycles in the absence of agricultural intensification when the sensitivity to resource deficiency is high (*σ* = 10). Solid lines are the average of 40 replications and the shading is the 95% confidence interval. b) Bifurcation diagram for the sensitivity to resource deficiency in the absence of agricultural intensification. When the sensitivity to resource deficiency is low the long-term state of the system is a sustainable steady state. When it is high the social-ecological system goes into collapse-recovery cycles. The purple lines depict the maximum and minimum values of the population density and the fraction of natural land during the oscillations. Solid lines are the average of 10 replications and the shading is the 95% confidence interval. c) Irreversible collapse dynamics of the social-ecological system in a scenario of low preference for intensification (*α* = 0.2). The sensitivity to resource deficiency is the same as in subfigure a). Solid lines are the average of 40 replications and the shades are the 95% confidence interval. d) Two-dimensional bifurcation diagram showing the different types of social-ecological equilibria as a function of the preference for intensification and the sensitivity to resource deficiency. The cycles of collapse and recovery (purple line) only exist in the absence of intensification. When the landscape is pristine, a greater preference for intensification increases the system’s resilience to increases in the sensitivity to resource deficiency. The frontier between the different equilibria was estimated by taking the average of the two closest points from one side and the other of the frontier. Each point is the average of 10 simulation replications.

An important factor determining whether the social-ecological system collapses is the speed of human-driven land use changes relative to the speed of demographic processes, controlled by *σ*, the sensitivity to resource deficiency. If the sensitivity to resource deficiency is low, human population density adjusts to a lower resource level faster, on average, than agriculture expands to increase resource production. Constraining the growth of population density has a stabilizing effect on the system dynamics and leads to a sustainable state in the long-term (Figure 1b blue line). When the sensitivity to resource deficiency is high, resource scarcity is compensated by agricultural expansion, leading to sustained growth of both agricultural land and population density that precludes stability within the system. As a consequence, the social-ecological dynamics enter cycles of reversible collapses (Figure 1a & Figure 1b purple lines).

Introducing agricultural intensification results in the disappearance of reversible collapses (Figure 1d). A small area of intense agriculture is sufficient to prevent landscape recovery following a collapse, making the degradation irreversible (Figure 1c). In contrast, a bias (i.e., high preference) for intense agriculture results in greater resilience against increases in resource deficiency sensitivity (Figure 1d). In this case, intensification limits agricultural expansion and decouples resource production from ecosystem services provision which contributes to stabilizing the system in a sustainable state (i.e., no percolation transition and *N* > 0.6).

### Collapse dynamics: percolation transition and habitat fragmentation

The abruptness of collapse is in sharp contrast to the initial phase of gradual land conversion (Figure 1a & Figure 1c). This sudden shift in the social-ecological dynamics is explained by the natural habitat undergoing a percolation transition. In the absence of agricultural clustering, the percolation transition occurs when the fraction of natural land is close to 0.59 (Figure 2a).

**Figure 2:**
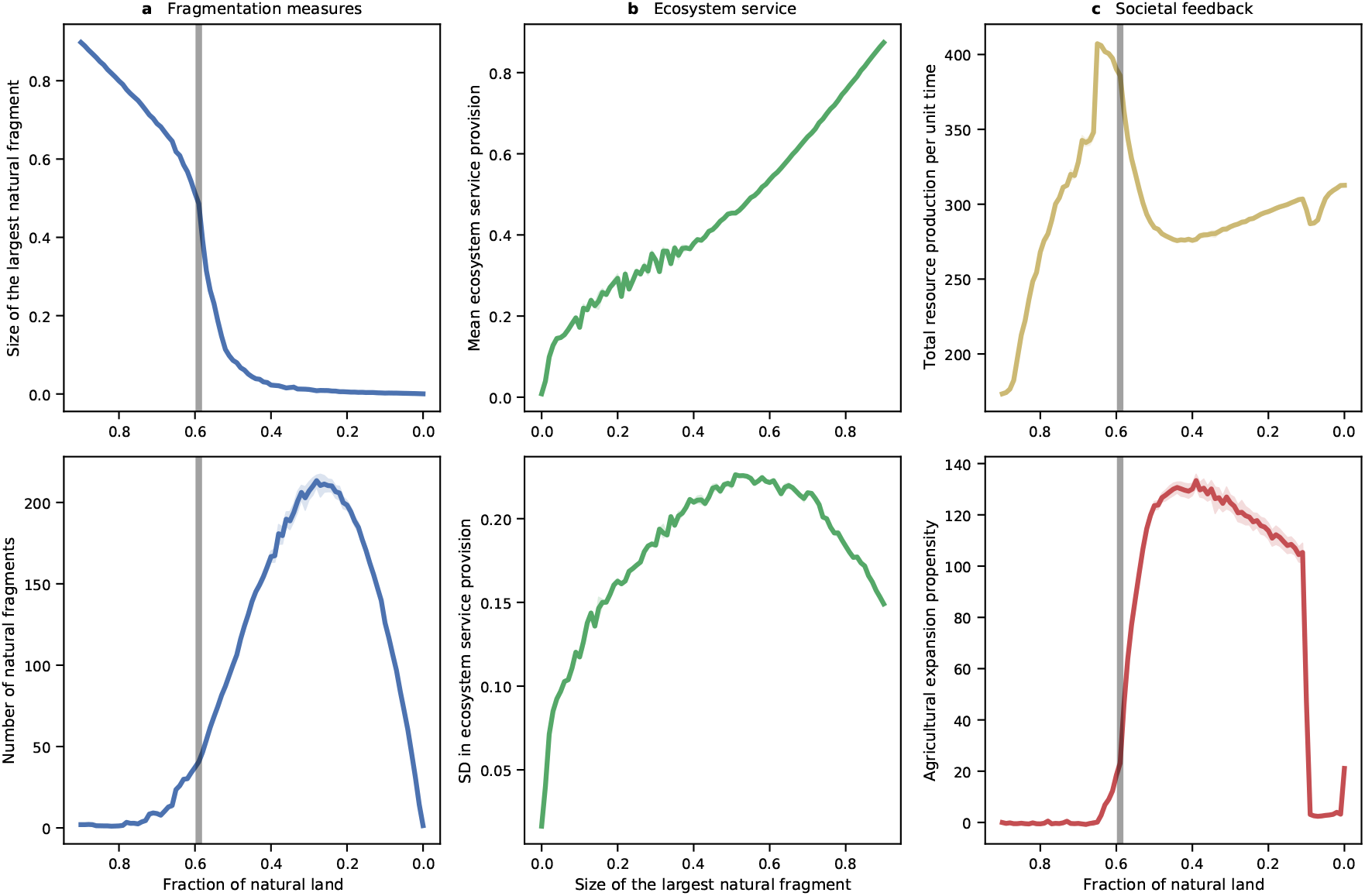
Signatures of a habitat percolation transition and the effects of habitat fragmentation on ecosystem services and land-use changes. a) Measures of habitat fragmentation. Both the size of the largest natural fragment in the landscape and the number of fragments show a highly non-linear relationship with the fraction of natural land. The abrupt decrease in the size of the largest natural fragments coincides with the abrupt increase in the number of fragments when the fraction of natural land reaches the percolation threshold (*N* ≃ 0.6, gray line in the plot). This is a signature of severe landscape fragmentation as a large number of small disconnected natural fragments emerge from the disappearance of a single large area of natural land. b) The average ecosystem services provision shows an almost linear dependence with the size of the largest natural fragment in the landscape. The spatial variance in ecosystem services provision increases as the size of the largest natural fragment decreases. The variance peaks when the size of the largest natural fragment is around half of the landscape size which coincides with the percolation transition (see panel a)). c) Agricultural production peaks just before the percolation transition and drops abruptly afterwards. This causes an explosive increase in the probability per unit time of agricultural expansion which results in rapid conversion to agriculture. Natural land conversion persists as long as agricultural production is below the desired level. The amount of cultivated area needed to satisfy resource demand increases with land conversion since habitat loss and fragmentation cause a systematic decrease in ecosystem services provision, thereby in agricultural productivity. The data for each curve comes from the realized dynamics of 20 model replications and the shades depict the 95% confidence interval.

The destruction of a small amount of habitat close to the percolation threshold (*N* ≃ 0.6) causes abrupt landscape fragmentation which is manifested in the sudden disappearance of a large natural landscape together with an increase in the number of disconnected natural patches (Figure 2a). The disappearance of large natural fragments results in a diminution of ecosystem services provision (Figure 2b) which translates into a marked reduction in resource production (Figure 2c). As a consequence, we observe a steep increase in the probability per unit time of expanding agriculture that leads to further habitat loss and fragmentation and worsens ecosystem services provision and therefore agricultural productivity. The social-ecological system is trapped in a positive feedback loop, where natural land is depleted to compensate for production losses, without success, more land is converted and the cycle continues.

### Preventing landscape fragmentation by clustering agricultural land

Agricultural clustering decouples habitat loss from habitat fragmentation, thereby diminishing the effects of a percolation transition (Figure 3a). With adequate clustering the risk of a percolation transition can disappear all together. When agricultural clustering is large, the linear relationship between the size of the largest natural fragment and the fraction of natural land reveals that natural habitat connectivity is preserved upon habitat loss (Figure 3a). In Figure 3b we depict the temporal changes of the fragmentation metrics to show that the percolation transition is avoided if agricultural clustering is high.

**Figure 3:**
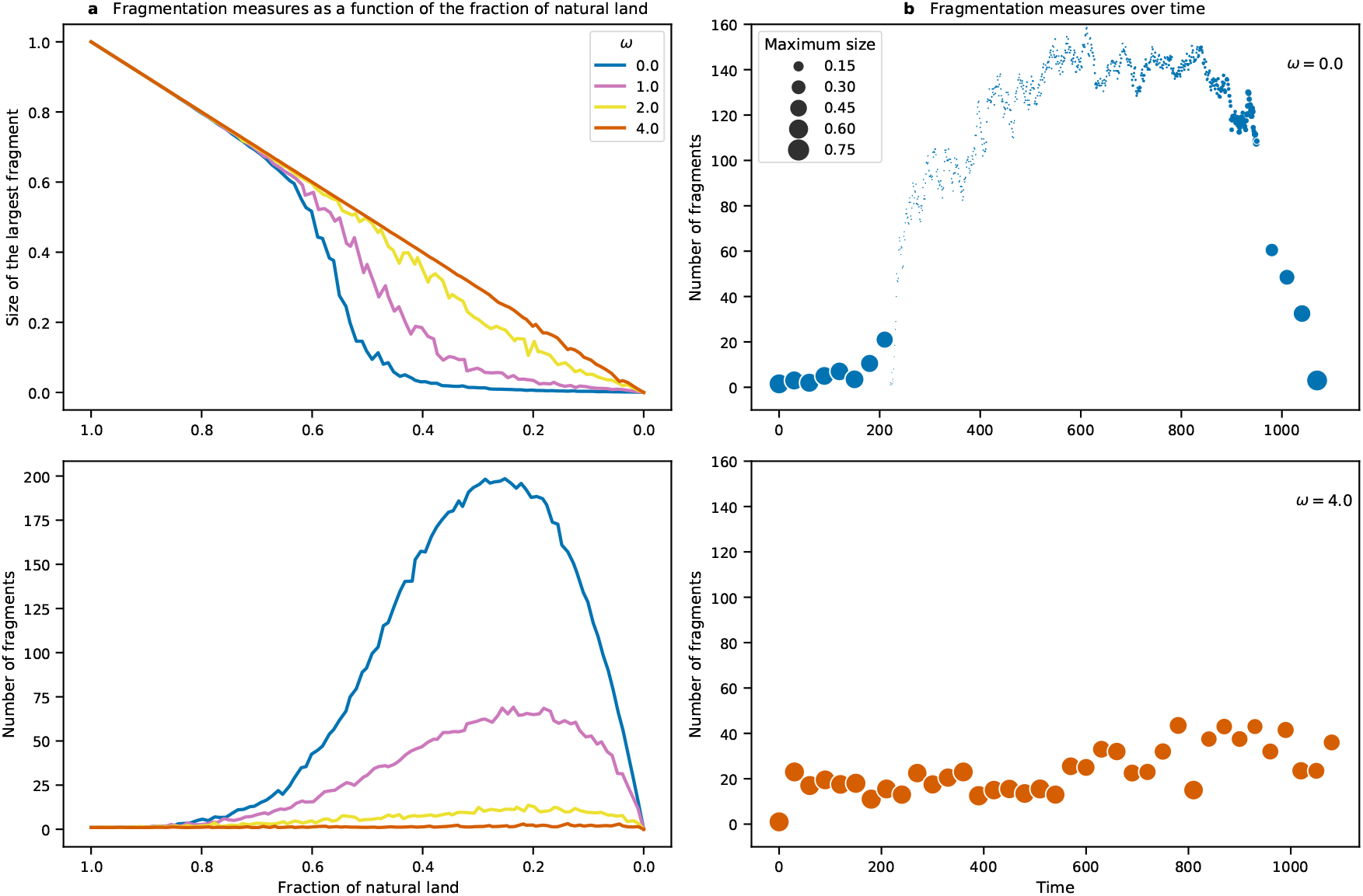
The effect of agricultural clustering on preserving landscape connectivity and the avoidance of a percolation transition. a) Fragmentation metrics as a function of the fraction of natural land for different values of agricultural clustering. Increasing agricultural clustering results in a linearization of the relationship between the size of the largest natural fragment and the fraction of natural land. This means that there is a decoupling between habitat loss and habitat fragmentation which results in the maintenance of natural land connectivity and the avoidance of a percolation transition. There are fewer disconnected natural patches as agricultural clustering increases, thus maintaining natural land connectivity. Each curve is the average of 100 simulations. b) Temporal changes in fragmentation metrics over time for no-clustering (top) and high-clustering (bottom). In the absence of agricultural clustering the changes in the size of the largest fragment and the amount of fragments are surprisingly abrupt. The time needed for landscape connectivity recovery is much greater than the time to widespread fragmentation. When agricultural clustering is high the habitat percolation transition is avoided and both the size of the natural fragments and the number of fragments are preserved over time. Each data point is the average of 40 simulations.

### Sustainable land-use management as a function of the landscape state

With the initial conditions set at low population densities in an almost pristine landscape, both the introduction of intensification and clustering have a stabilizing effect on the system, generating dynamics that evolve to sustainable steady-states in the long-term (Figure 1d & Figure 3a). An analysis of the effect of the landscape’s initial conditions on different management strategies showed that their success is highly dependent on the initial fraction of natural land (Figure 4). Greater preference for intense agriculture reduces the range under which sustainability is reached. We observe that neither land sharing (low intensification, low clustering) nor land sparing (high intensification, high clustering) are the most resilient management strategies, but rather a combination of high clustering and low intensification (Figure 4).

**Figure 4:**
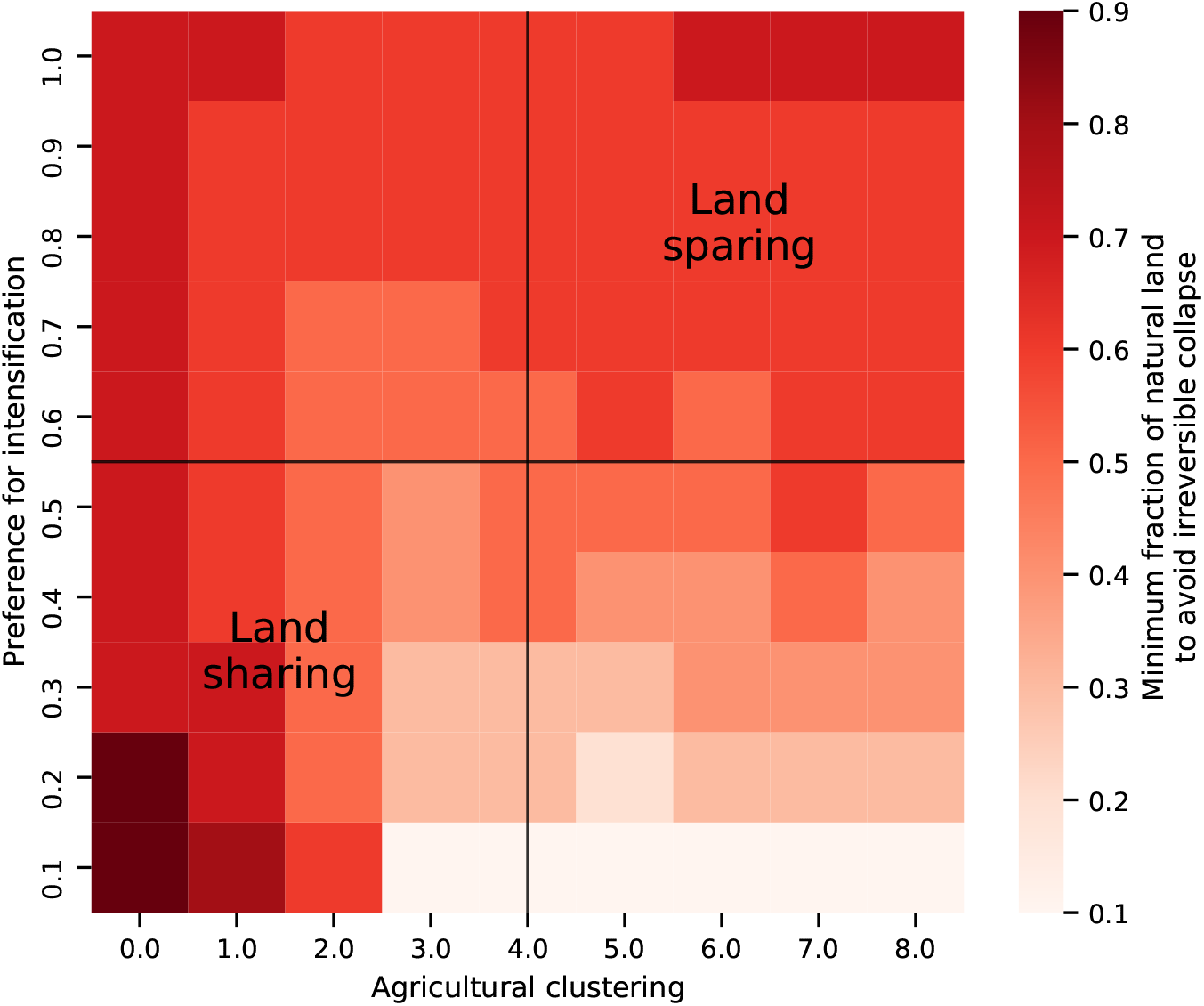
Suitability of different land-use management strategies as a function of the landscape’s initial conditions. The colors show the minimum fraction of natural land needed to be certain that a given land-use management strategy will not lead to an irreversible collapse. This puts into perspective some of the aforementioned results, as it shows that the success of intensification strategies are highly dependent on the amount of natural land in the landscape when intensification occurs. The figure shows that agricultural clustering is effective in increasing sustainability but that intensification reduces its efficiency (Land sparing square). More interestingly this suggests that neither land sharing nor land sparing are the most suitable strategies towards sustainability. Each point is the average of 10 simulations.

## 4 Discussion

Among the seventeen United Nations’ Sustainable Development Goals, two are related to sustainable resource production and food security and another three to environmental conservation issues (UN, 2015). Agricultural management lies at the heart of the sustainability debate. Agricultural expansion and intensification have greatly increased food availability, keeping pace with growing resource demands in recent decades (Barrett *et al*., 2020). But these practices also deeply transform landscapes across the world and, by doing so, drive the loss and degradation of natural habitats (Kehoe *et al*., 2017). This has prompted society to question whether it is possible to promote food security and environmental conservation. We demonstrated how gradual habitat loss can cause abrupt habitat fragmentation that impairs ecosystem services provision, thus resulting in a large reduction of agricultural production. We also showed how naive and uninformed management responses to the production drop (i.e., continuous conversion to agriculture without gains in yield) deepen landscape degradation and fail to compensate production losses. Long-term dynamics in the model show either a collapse or continued cycles. These practices are unsuccessful at both securing food resources and conserving the environment.

We approached land use management from an social-ecological perspective, modeling the likelihood of agricultural expansion and intensification as a function of the population’s perceived lack of resources. From this perspective, we showed that the main factor determining whether the social-ecological system collapses is the speed of human-driven land use changes relative to the speed of demographic processes. When the population’s reaction to a lack of resources is slow compared to demographic timescales, population numbers adjust to resource availability before agriculture is expanded or intensified in order to increase production. The social response to an insufficient level of resources is, in this case, a reduction of the population’s pressure on the environment. On the contrary, when there is an urgent response to a perceived lack of resources compared to demographic timescales, small production drops are systematically replenished by further agricultural expansion and intensification. In this scenario, the population’s response is to increase pressure on the environment which can result in a social-ecological collapse.

Reducing per capita consumption in response to a lack of resources is an alternative to reducing population numbers, and is equally effective in terms of decreasing the human pressure on the environment. Due to its simplified nature, our model does not account for changes in consumption levels as a reaction to changes in resource availability. Modifying the model to account for these changes, however, would not affect our main findings since the effects of per capita consumption levels and population size on the environment are the same, and one variable can be substituted for the other. Our results stress that the path to sustainability relies on the society’s ability to determine when an increasing societal pressure on the environment can lead to severe, and possibly irreversible, environmental degradation.

We illustrated how systematic and progressive natural habitat depletion as a response to insufficient resource availability causes the landscape to undergo a percolation transition (Aharony & Stauffer,2003; Taubert *et al*., 2018). Once a threshold in the fraction of natural habitat is attained, the conversion of a small number of natural patches to agriculture causes the abrupt fragmentation of a single large natural cluster. This results in the proliferation of many small disconnected natural patches and the dissipation of large patches. We found that such major fragmentation events resulted in a prominent decrease in ecosystem service provision. The amplitude of the decrease depends on how ecosystem service supply scales with the area of natural fragments (Mitchell *et al*., 2014, 2015a). The model describes clustering as touching, connected neighboring patches, but in reality ecosystem functioning may not require direct connectivity between natural patches. We did not broaden our analysis to include how ecosystem services scale with area or how connectivity varies with the distance between patches. The estimation of both quantities is most likely dependent on both the ecosystem and the services one considers. However, empirical evidence on the negative effects of fragmentation on ecosystem functioning supports the theoretical claim that a landscape percolation transition would greatly affect ecosystem services provision (Haddad *et al*., 2015).

In our model, the large drop in ecosystem service provision after the percolation transition causes a significant decrease in resource production. As a consequence, agricultural expansion accelerates, and further aggravates habitat loss and fragmentation and the subsequent reductions in ecosystem service provision and agricultural productivity. Agricultural expansion becomes less efficient over time and can result in a vicious cycle, if unchecked. Increasing pressure on the environment close to the percolation threshold triggers a social-ecological collapse driven by naive and uninformed responses to decreases in ecosystem service provision.

The amplitude of the collapse depends on the degree of crop dependency on ecosystem services. A thorough quantitative estimation of the contribution of different ecosystem services to agricultural production is currently lacking. Substantial progress has nevertheless been made on the quantification of some essential services, such as pest control (Dainese *et al*., 2019), pollination (Garibaldi *et al*., 2011), water quantity and quality (Swift *et al*., 2004) and soil structure and fertility (Tsiafouli *et al*., 2015). The weight of these services on production varies with the type of crop, geographical regions and the type of agriculture (Gallai *et al*., 2009). The decline in pollinators has raised food supply concerns, on a global-scale, as approximately 40% of total food crop production comes from animal pollinated crops (Power, 2010). Our model does not take into account alternative crops. For example, cereal production does not depend on animal pollination, therefore population collapses that emerge from our model could be avoided by altering crops rather than simply turning to expansion or intensification. However, this is just one ecosystem service. Changing crops to non-pollinated varieties would result in greatly simplified diets and nutrient deficiencies, as fruit and vegetable production would drastically decline (Gallai *et al*.,2009). These crops would also be vulnerable to a decline in other ecosystem services. Countries in the Global South are in general more vulnerable to decreases in ecosystem services provision because they lack assistance from technology and institutional support that buffer against ecosystem service loss. Overconsumption in other countries, primarily in the North, threaten ecosystems in the South, making this a complex social-ecological, agricultural and economic issue.

Agricultural intensification, in the sense of increasing productivity per unit area, has been proposed as a solution to meet future food demands while sparing natural land from conversion (Phalan *et al*., 2011). By replacing ecosystem services with synthetic inputs, intensification decouples agricultural productivity from the environment to some extent. As a result agricultural yields and their stability are increased, at least in the short term. However, intensive agriculture takes a greater toll on the environment in other ways (Tilman, 1999; Chappell & LaValle, 2011; Tscharntke *et al*., 2012), which can have long-term repercussions. Pollution and eutrophication of groundwater systems are a common consequence of agricultural intensification due to the continuous application of fertilizers and pesticides that saturate in the soil (Dale & Polasky, 2007; Galloway *et al*., 2008). Overcropping to increase yields also affects soil structure and natural vegetation loss leads to soil erosion (Montgomery, 2007). Intensification can also affect yield stability; for example, intensive agriculture reduces the capacity of agricultural systems to buffer climatic hazards.

It is also unclear whether intensification truly promotes land sparing (Tscharntke *et al*., 2012; Kremen, 2015). Empirically a correlation between agricultural contraction and intensification has not been observed (Kremen, 2015), probably for social-economic reasons. The gains in efficiency that are brought by intensification increase economic profit and thus fuel the expansion of intensive agriculture (Gusso *et al*., 2017). Our results highlight that intensification can be a successful strategy towards sustainability provided it is accompanied by real conservation of natural habitats. However, if conservation is ignored, we showed that the increase in degradation caused by slight increases in agricultural intensification can lead to social-ecological collapses. Furthermore, we showed that a land sparing strategy was not optimal in landscapes without large amounts of natural land. Present work on agro-ecological intensification promotes crop productivity through ecosystem services management, which has the potential to maintain spared natural landscapes by reducing environmental degradation from synthetic inputs (Bommarco *et al*., 2013; Garnett *et al*., 2013).

Green (2005) seminal work on wildlife-friendly-farming and land sparing provided a framework for research about the best land-use management strategies that jointly achieve sufficient resource production and conservation of the environment (Green, 2005). Our work adds crucial components to the debate, i.e. landscape structure and connectivity of natural fragments. It shows that clustering agricultural land together has the positive effect of preserving large natural fragments that maintain ecosystem services in the landscape. When agriculture is uniformly distributed in space, the risk of agricultural expansion causing a percolation transition is higher. However, our results stress that successful management strategies are highly dependent on the state of the landscape as landscapes with highly clustered intense agriculture need very large fractions of natural land to be sustainable in the long term. This puts the land sharing vs. land sparing debate into perspective, as high clustering combined with low-intensity agriculture emerged as the most sustainable strategy in highly managed landscapes.

Landscapes around the world are constantly changing to accommodate changes in human population and societies’ demands. Our work shows how abrupt transitions in landscape structure can occur through gradual landscape changes. Since estimating minimum areas to provide some services can be an almost impossible task and is highly region-dependent, our work stresses the need for precautionary behavior in designing management policies and suggests preserving large natural fragments in complex landscapes as a preventive measure. It is important to manage agricultural landscapes in a way that preserves connectivity of natural fragments to avoid a possible collapse in ecosystem service provision.

Globalization has displaced croplands to most pristine regions of the world, decoupling production from consumption. This decoupling could delay or mask human-nature feedbacks, but it does not diminish the importance of sustainable agricultural management strategies. Our work clearly shows that heterogeneous (non-clustered) intensive agriculture can lead to social-ecological collapses by way of percolation transitions; decision-making processes and perceptions within society will determine whether the percolation threshold will be crossed. Further studies focusing on the interactions between management practices, cultural traits and economics are encouraged to identify possible strategies that aim to conserve natural land in combination with intensification. Greater level of detail on the crops and ecosystem processes to be considered is needed for precise guidance. Lastly, the development of models to study the coupling of different landscapes and human populations through migration and trade would be an important step to better understand the conditions under which sustainability can be attained.

## Acknowledgements

This work was supported by the TULIP Laboratory of Excellence (ANR-10-LABX-41) and was conducted within the framework of the BIOSTASES Advanced Grant funded by the European Research Council under the European Unions Horizon 2020 research and innovation programme (grant agreement no. 666971).

## Authorship

All authors contributed to the design of the study. DBP elaborated the model, performed and analyzed the simulations, and wrote the first draft of the mansucript. All authors contributed substantially to revision.

## Data accesibility statement

Data available from the repository https://doi.org/10.5281/zenodo.4905944.

