## Supplementary Material for "Habitat percolation transition undermines sustainability in social-ecological agricultural systems"

June 7, 2021

#### Appendix 1 : Derivation and non-dimensionalisation of model's equations

Here we follow the structure of the Model and Methods section of the main text and introduce greater details into the derivation of the equations and its non-dimensionalisation.

##### Ecosystem services provision

As explained in the main text, we formalize ecosystem services as a quantity generated by natural fragments and flowing to the rest of the landscape. We assume that the magnitude of ecosystem services generated by a fragment saturates with its area and choose to use a powar-law function with exponent smaller than 1 to model the saturation. We further assume that the services flow from a natural patches to their closest neighbours defined in this model as the Von-Neumman neighbourhood in the lattice, i.e. the northern, eastern, southern and western cells. The amount of ecosystem services $E_i$ that nature provides to landscape cell $i$ is:

$$E_i = \sum_{j \in N_i} E_0 \, A_j^z, \quad (1)$$

where $N_i$ is the set of natural neighbours of cell $i$, $E_0$ is the prefactor of the saturation function, $A_j$ is the area of natural neighbour $j$ of cell $i$ and $z$ is the saturation exponent. Given a landscape size, a cell receives the maximum possbile amount of ecosystem services when the whole landscape is in a natural state. The maximum possible amount of ecosystem services provided to a cell $E_{max}$ can then be written:

$$E_{max} = 4 \, E_0 \, A_L^z, \quad (2)$$

where $A_L$ is the total area of the landscape and prefactor 4 comes from the sum of the contribution over the 4 neighbours. We non-dimensionalize the measure of ecosystem service provision by expressing as a fraction of the possible maximum. The non-dimensional ecosystem service provision $\epsilon$ for cell $i$ is written:

$$\epsilon_i = \frac{E_i}{E_{max}} = \frac{1}{4} \sum_{j \in N_i} \left( \frac{A_j}{A_L} \right)^z = \frac{1}{4} \sum_{j \in N_i} a_j^z, \quad (3)$$

where $a_j = A_j / A_L$ is the fraction of the landscape occupied by natural fragment $j$.

#### Production of agricultural resources

Total agricultural production per unit time in the landscape is obtained by summing over the production per unit time of every low-intense and high-intense agricultural cells. We assume that the low-intense agricultural production is possitively affected by the amount of ecosystem services the agricultural cell receives, hence we write it as the sum of a baseline production and a contribution related to ecosystem service provision. On the contrary, we assume that resource production in high-intensity cells is fixed and completely independent of ecosystem service provision. We write total resource production $\mathcal{Y}$ as follows:

$$\mathcal{Y} = \sum_{i \in A_l} (\mathcal{Y}_0 + Y_E E_i) + \sum_{i \in A_h} \mathcal{Y}_1, \quad (4)$$

where $A_l$ and $A_h$ are the sets of low-intense and high-intense agricultural cells respectively, $\mathcal{Y}_0$ is the baseline production per unit time of low-intense agricultural cells, $Y_E$ is the production per unit time per unit of received ecosystem service of low-intense agricultural cells and $\mathcal{Y}_1$ is the production per unit time of high-intensity agricultural cells. We non-dimensionalize total resource production by expressing it relative to the maximum possible contribution of ecosystem services to resource production: $Y_E E_{max}$. This yields the following expression for non-dimensional total resource production per unit time $Y$:

$$Y = \frac{\mathcal{Y}}{Y_E E_{max}} = \sum_{i \in A_l} \left( \frac{\mathcal{Y}_0}{Y_E E_{max}} + \epsilon_i \right) + \sum_{i \in A_h} \frac{\mathcal{Y}_1}{Y_E E_{max}} = \sum_{i \in A_l} (y_0 + \epsilon_i) + \sum_{i \in A_h} y_1, \tag{5}$$

where $y_0$ and $y_1$ are non-dimensional parameters that represent the baseline productions per unit time of low-intense and high-intense agriculture relative to the ecosystem services contribution respectively.

#### Differential equation for Human demography

We assume that Human population dynamics follows logistic growth with a time-evoloving carrying capacity determined by the maximum population density that can be sustained given an amount of resource production $\mathcal{Y}$ and a per capita consumption $c$ per unit time. The equation for Human population density $P$ is then as follows:

$$\frac{dP}{dT} = rP \left( 1 - \frac{cP}{\mathcal{Y}} \right), \tag{6}$$

where $r$ is the growth rate of the population density when density is very low compared to the carrying capacity $\mathcal{Y}/c$. We choose to non-dimensionalize the population density by expressing it relative to the population density that can be sustained via the maximum possible contribution of ecosystem services in one low-intensity agricultural cell $Y_E E_{max}$. This means that the non-dimensional population density $p$ can be written $p = c\,P/(Y_E E_{max})$ and By replacing in equation 6 and non-dimensionalizing the time by expressing it relative to the inverse of the population growth rate $r$, we obtain the following non-dimensional equation for the Human demography:

$$\frac{dp}{dt} = p \left( 1 - \frac{p}{Y} \right). \tag{7}$$

### Propensities of land use/cover transitions

We derive the propensities of land use/cover transitions by defining the average time that it takes a given transition to occur given the system's state. Assuming that land use/cover transitions are a memory-less stochastic process, the propensity, here defined as the probability per unit time, of a transition can be written as the inverse of the expected time that one should wait to observe the transition given the state of the system. Hereafter we detail the reasoning behind the equations for the average expected waiting times before the transitions and show the obtention of the propensities' expressions.

##### Agricultural expansion and intensification

The expected time $T_A$ before expansion or intensification of agriculture depends on the resource deficit $\Delta R = cP - \mathcal{Y}$ experienced by the human population. The greater the resource deficit, the faster Humans respond by enlarging agricultural activities via agricultural expansion or intensification. We further assume that this time tends to the infinity as the resource deficit tends to zero, and that it tends to zero as the resource deficit tends to the infinity. If the resource deficit is negative, meaning that resource production exceeds the demand neither expansion nor intensification can occur, hence $T_A \rightarrow \inf$ when $\Delta R < 0$. Hereafter we will only write the equations in the case that $\Delta R > 0$. A parsimonious choice for a function satisfying such demands is an inverse function of the resource deficit:

$$T_A = \tau_A \frac{1}{\Delta R} = \tau_A \frac{1}{cP - \mathcal{Y}}, \tag{8}$$

where $\tau_A$ is rate of decrease of the average time before the transition per unit of resource deficit. Re-scaling the time by $1/r$ and replacing population density and total resource production by their expression in function of their non-dimensional equivalents we obtaint the following non-dimensional equation for the average aiting time before agricultural expansion or intensification:

$$rT_A = r\tau_A \frac{1}{\Delta R} = \frac{r\tau_A}{Y_E E_{max}} \frac{1}{p - Y}, \tag{9}$$

The non-dimensional propensity $\pi_A$ of agricultural expansion or intensification occurring somewhere in the landscape is then obtained by taking the inverse of equation 9:

$$\pi_A = \frac{Y_E E_{max}}{r \tau_A} (p - Y) = \sigma \Delta r, \tag{10}$$

where $\sigma$ (non-dimensional) is the sensitivity to resource deficit and represents the increase rate of epxnasion or intensification propensity per unit of resource deficit, and $\Delta r$ is the non-dimensional resource deficit.

We assume that whether expansion or intensification is chosen depends on the value of parameter $\alpha$ which represents the preference for agricultural intensification and is the probability of choosing intensification over expansion if an agricultural land use transition is to occur. The expansion $\pi_E(i)$ and intensification $\pi_I(i)$ propensities in cell $i0$ assuming that both processes occur with uniform probability in space are then :

$$\pi_E(i) = \begin{cases} \frac{1}{N} \sigma \, \Delta r \, (1 - \alpha), & \text{if } \Delta r > 0 \\ 0, & \text{otherwise.} \end{cases} \tag{11}$$

$$\pi_I(i) = \begin{cases} \frac{1}{A_l} \sigma \, \Delta r \, \alpha, & \text{if } \Delta r > 0 \\ 0, & \text{otherwise.} \end{cases} \tag{12}$$

To study the influence of non-uniform spatial configurations of agriculture in the landscape on the social-ecological dynamics we introduce an agricultural clustering parameter $\omega$ which controls the likelihood of converting cells that are neighbours of other cells with the same type of land cover after the transition. For example, if $\omega > 0$ it is more likely to convert to agriculture a natural cell that is in the neighbourhood of a low-intensity agricultural cell than a natural cell surrounded by other natural cells, or high-intensitiy cells. The same applies to intensification: low-intensity agricultural cells close to high-intensity ones would be prefered. As $\omega$ increases the probability of converting cells close to already converted ones increases and the distribution of agricultural land becomes increasingly clustered by land use type. The dependence of the expansion or intensification propensities with respect to the clustering parameter $\omega$ is:

$$\pi_E(i) = \begin{cases} \frac{1}{\mathcal{N}}\,\sigma\,\Delta r\,(1-\alpha)\ [\max(0.1, A_l(i))]^{\omega}\,, & \text{if } \Delta r > 0 \\ 0, & \text{otherwise.} \end{cases} \tag{13}$$

$$\pi_I(i) = \begin{cases} \frac{1}{\mathcal{N}}\,\sigma\,\Delta r\,\alpha\ [\max(0.1, A_h(i))]^{\omega}\,, & \text{if } \Delta r > 0 \\ 0, & \text{otherwise.} \end{cases} \tag{14}$$

$$\tag{15}$$

where $A_l(i)$ and $A_h(i)$ are the set of low-intesity and high-intensity agricultural neighbours of cell $i$ respectively. This formulation means that the likelihood of choosing a cell in the neighborhood of other cells of the same desired kind is $(10\#\text{Neighbours})^{\omega}$ larger compared to a cell without any neighbour of the desired kind. The normalization factor $\mathcal{N}$ guarantees that the total propensity of intensification or expansion to occur is always $\sigma\,\Delta r$:

$$\pi_A = \sum_{i \in N} \pi_E(i) + \sum_{i \in A_l} \pi_I(i) = \sigma\,\Delta r, \tag{16}$$

where $N$ and $A_l$ are the sets of natural and low-intensity agricultural cells of the landscape respectively.

##### Fertility loss of agricultural land

We model the lost of agricultural cells are lost as a consequence of fertility loss (Pimentel, 2006) and assume that the time until fertility loss $T_L(i)$ of an agricultural cell $i$ depends on the amount of ecosystem services that it perceives. We further assume that $T_L(i)$ tends to the infinity as $E_i$ tends to $E_{max}$ and that $T_L(i)$ is finite and positive when $E_i = 0$. A parsimonious choice for a function satisfying the previous demands is:

$$T_L(i) = \tau_L \frac{1}{E_{max} - E_i}, \tag{17}$$

where $\tau_L$ is the rate of increase of the average fertility loss time per unit of ecosystem service provision. AS previosuly, we re-scale the times by $1/r$, and introduce the non-dimensional ecosystem services provision $\epsilon_i$ in the previosyu equation to obtain the following non-dimensional equation:

$$rT_L(i) = \frac{r\tau_L}{E_{max}} \frac{1}{1 - \epsilon_i}, \tag{18}$$

Taking the inverse of the previous equation yields the following expression for the non-dimensional fertility loss propensity $\pi_L(i)$ in cell $i$:

$$\pi_L(i) = \frac{E_{max}}{r\tau_L} (1 - \epsilon_i) = \rho_L (1 - \epsilon_i), \tag{19}$$

where $\rho_L$ (non-dimensional) is fertility loss's sensitivty to ecosystem service provision and represents the decrease rate of fertility loss propensity per unit of ecosystem service. The fertility loss process is identical for low-intensity and high-intensity agricultural cells, with the exception that low-intensity agricultural cells transition to a natural state and high-intensity ones transition to a degraded state when they lose fertility. The assumption behind this modelling choice, is that the time-scales at which an old low-intensity agriculutral field goes back to a natural state is fast relative to other land cover transitions, hence we make the approximation of assuming that it happens instantaneously in the model.

**Spontaneous land recovery and degradation**

We also model the average time until land recovery $T_R$ or degradation $T_D$ in cell $i$ as a function of the ecosystem services $E_i$ that the cell receives. We assume that the dependence of land degradation average time on ecosystem service provision is identical as the one of fertility loss. Following the same steps as before we arrive to the following non-dimensional expression for the land degradation propensity $\pi_D(i)$ in cell $i$:

$$\pi_D(i) = \rho_D (1 - \epsilon_i), \tag{20}$$

where $\rho_D$ (non-dimensional) is land degradation's sensitivity to ecosystem service provision and represents decrease rate of land degradation propensity per unit of ecosystem service.

For land recovery, we assume that $T_R(i)$ tends to the infinity when $E_i$ tends to 0 and that it has a finite and positive value when $E_i = E_{max}$. A parsimonious choice for a function satisfying the presvious demand is:

$$T_R(i) = \tau_R \frac{1}{E_i}, \tag{21}$$

where $\tau_R$ is the rate of decrease of the average land recovery time per unit of ecosystem service. By introducing the non-dimensional from of $E_i$ and re-scaling the time by $1/r$ we obtain the following non-dimensional equation:

$$rT_R(i) = \frac{r\tau_R}{E_{max}} \frac{1}{\epsilon_i}, \tag{22}$$

and taking the inverse we obtain the expression of the non-dimensional land recovery propensity $\pi_R(i)$ in cell $i$ :

$$\pi_R(i) = \rho_R \epsilon_i, \tag{23}$$

where $\rho_R$ is land recovery's sensitivity to ecosystem service provision and it represents the increase rate of land recovery propensity per unit of ecosystem service.

### Appendix 2: Estimation of parameter values

We searched the litterature for orders of magnitude caracterising the different processes we incorporated in the model to estimate the parameters' values. This approach yields parameter values that are not precise enough to provide highly confident quantitative predictions of future land-use and population changes. It does however set reasonable scales for their value that allow us to explore different possible scenarios for the future and provide plausible qualitative predictions as well as mechanistic explanations of the processes leading to habitat loss and fragmentation that emerge from the bi-directional feedbacks between Humans and Nature. This approach is justified by the extreme difficulty of making precise estimations of this kind of paramters from empirical data. For example, data on the timescales of the recovery of degraded land is highly dependent on geographical locations and considered ecosystems and quantifications of the relationship between the level of ecosystem service provision and the speed of recovery do not exist to our knowledge. What we can do to surpass this issue is to use paramter values that are in the order of magnitude of such processes. Below we detail the reasoning and provide the references that we used to estimate parameter values. We do not discuss neither the estimation of the per capita resource consumption or of agricultural yields as they only intervene in the non-dimensionalization of Human population density and do not have a qualitative influence in the dynamics of the non-dimensional model. The only information that can be gathered by their estimation is a quantitative prediction of changes in Human population density but that is not the goal of our study.

#### Estimation of the saturation exponent of ecosystem service provision with area $z$

The saturation exponent of ecosystem services provision with the area of natural fragments was set to $z = 1/4$ so that a ten-fold decrease in a fragment's area causes around of a two-fold decrease of ecosystem service provision. Decreasing the saturation exponent (i.e. increasing saturation) has the effect of containing the propagation of degraded land but enlargens variability in the dynamics (Figure 1). It also contains the negative effects of fragmentation in the wake of the percolation transition (Figure 2c) since large ecosystem service provision is maintained for relatively small sized natural fragments (Figure 2b). On the contrary, increasing the saturation exponent makes the sytem more vulnerable to habitat loss and fragmenation, as ecosystem service provision decreases too fast with fragment area reaching almost a linear relationship between mean ecosystem service provision and the area of the largest natural fragment (Figure 2b). This has the effect of diminishing the likelihood of recoveries after social-ecological collapses (Figure 1).

#### Estimation of the low-intensity and high intensity baseline yields $y_0$ and $y_1$ (non-dimensional)

The baseline yield of low-intensity agriculture was estimated based on evidence from the litterature on the contribution of natural pollination and pest control on agricultural yields (Bommarco et al., 2013; ?). Extrapolating to account for the contribution of other non-accounted ecosystem services we defined that 4/5 of the production in low-intensity agricultural patches depends on ecosystem service provision. Decreasing the dependance on ecosystem service provision (i.e. increasing baseline production) contains agricultural expansion in the ake of the percolation transition (Figure 3), as less land is required for the same production

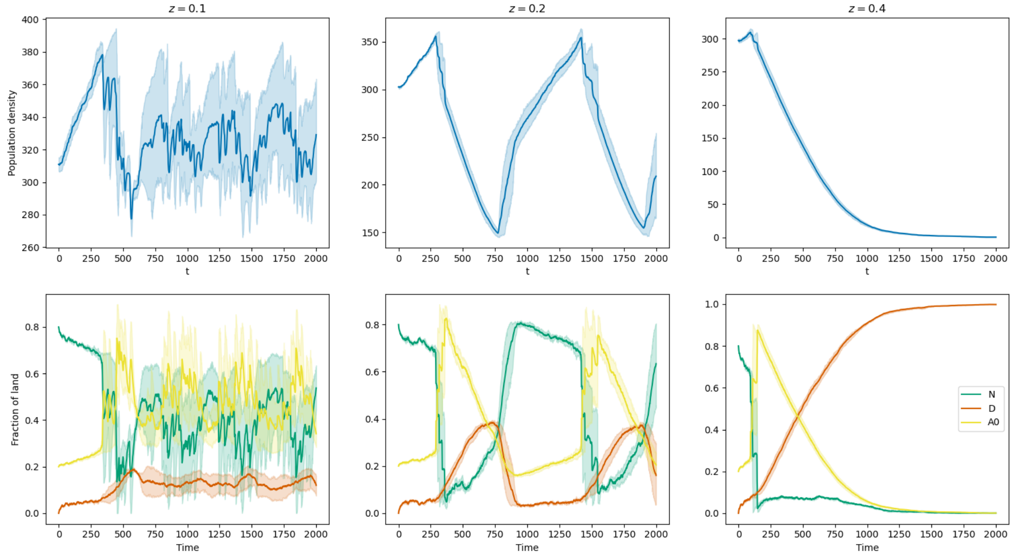


**Figure 1:** Social-ecological dynamics for different values of the ecosystem service saturation exponent. Solid lines are the mean of 5 replications and the shades represent the 95% confidence interval. Low values of the exponent (i.e. high saturation) help the containment of land degradation but dynamics become increasingly variable. As $z$ increases in value the amplitude of the collapses increases. High values of the exponent (i.e. low saturation) render collapses irreversible. In these simulations: $a = 0$, $w = 0$, $\sigma = 10$, $a_0 = 0.2$ and the rest of the parameters are set to the value presented on the main text table.

in the absence of ecosystem services (Figure 4). However, the effects of the percolation transition are not attenuated as increasing the decoupling between agricultural yields and ecosystem service does not help in containing land degradation. Precise assessements of the contribution of ecosystem services on agricultural are needed to make confident assessements of the sustainability of agricultural landscapes in the presence of habitat loss and fragmentation. The yield of high-intensity agricultural patches was then chosen so that it equals the yield of low-intensity patches when ecosystem service provision is at its possible maximum (Chappell and LaValle, 2011).

#### Estimation of the population growth rate $r$ (dimensional)

Parameter $r$ is Human density growth rate when the density is very low compared to the carrying capacity. A precise estimation of $r$ with empirical data would require data on the difference between per capita food supply and per capita resource consumption across time, as their ratio would be the Human carrying capacity

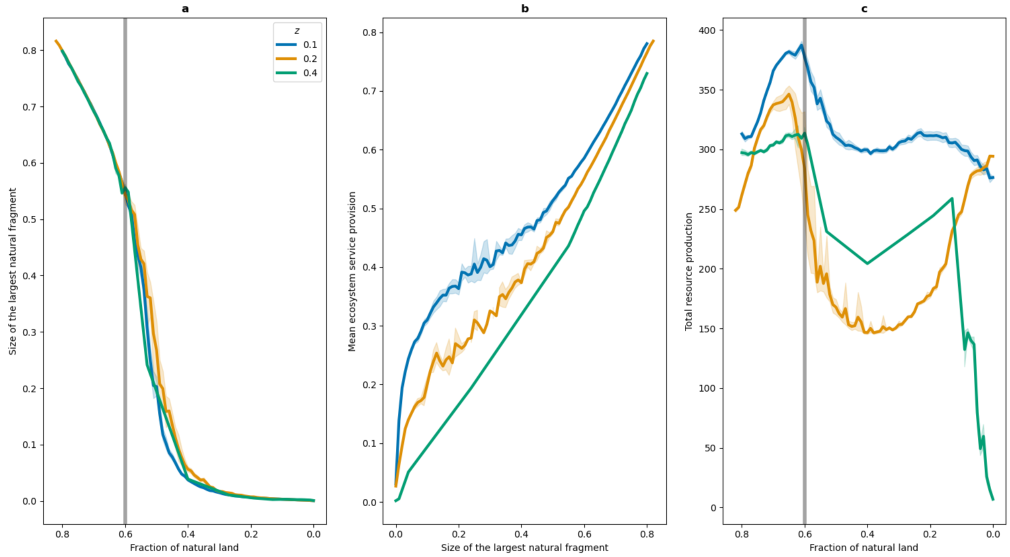


**Figure 2:** Landscape structure and consequences on ecosysetm service provision and agricultural production for different values of the ecosystem service saturation exponent. Solid lines are the mean of 5 replications and the shades represent the 95% confidence interval. a) Size of the largest natural fragment as a function of the fraction of natural land. The large decrease on the size of the largest fragment at the percolation transition is the same no matter the value of $z$. b) Lower values of the saturation exponent (i.e. high saturation) helps preserving ecosystem service provision when giant natural fragments dissapear. c) The amplitude of the production drop increases when the saturation exponent increases (i.e. lower saturation). In these simulations: $a = 0$, $w = 0$, $\sigma = 10$, $a_0 = 0.2$ and the rest of the parameters are set to the value presented on the main text table.

(which changes over time). Knowledge of the Human carrying capacity across time would allow to fit a logistic function to population trends and obtain an estimationof parameter $r$. In standard datasets (FAOSTAT for example) per capita consumption data is actually data on food supply making the distinction between both impossible. To overcome this limitation we estimate that population density in the presence of an almost infinite amount of resources (i.e. population density very low compared to the carrying capacity) can double in a time ranging from 1 year (human pregnancy timescale) to 1 decade. The doubling time at very low population densities compared to the carrying capacity is $\ln(2)\,1/r \simeq 1/r$. In our simulations we assumed $1/r = 2$ which gives a doubling time around of 2 years.

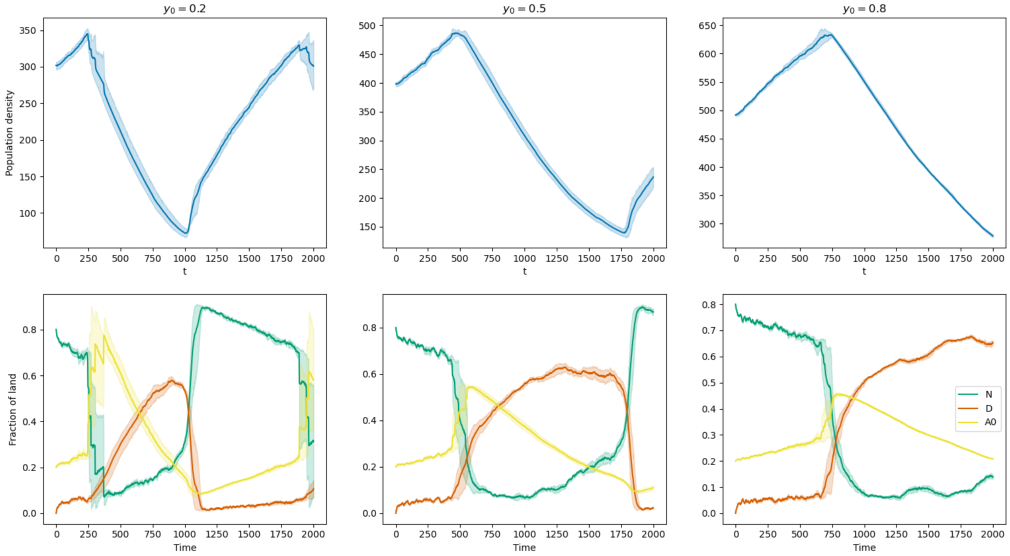


**Figure 3:** Social-ecological dynamics for different values of the baseline yield of low-intensity agriculture. Solid lines are the mean of 5 replications and the shades represent the 95% confidence interval. Increasing the baseline production decreases the dependency of agricultural production one cosysetm services and thus makes Humans response to the percolation transition less intense. Even thoufh agricultural expansion is accelerated, it doesn't need to explode as much to compensate production losses due to habitat fragmentation. This does not contain land degradation resulting in reversible collapses. The frequency of the collapse-recovery cycles decreases as the baseline production $y_0$ increases. In these simulations: $a = 0$, $w = 0$, $\sigma = 10$, $a_0 = 0.2$ and the rest of the parameters are set to the value presented on the main text table.

#### Estimation of land use transitions sensitivity to ecosystem service provision $\rho_R$, $\rho_D$, $\rho_L$ (non-dimensional)

We searched the litterature for references in the characteristic timescales of degraded land's recovery. Based on **??**, we made the estimation that degraded lands surrounded by good quality habitat can be recolonized in times of the order of the decade. Hence we assume land recovery's sensitivity to ecosystem services (non-dimensional) is $\rho_R \simeq 0.2$. This means that the average land recovery time is of 10 years when the ecosystem service provision is at the maximum possible level.

We placed ourselves in the optimistic scenario where in the absence of Humans a landscape composed by 10% of natural land and 90% of degraded land is able to fully recover in the long-term with probability equal to 1 (see Fig 5). A reasonable assumption for land degradation's sensitivity to ecosystem service

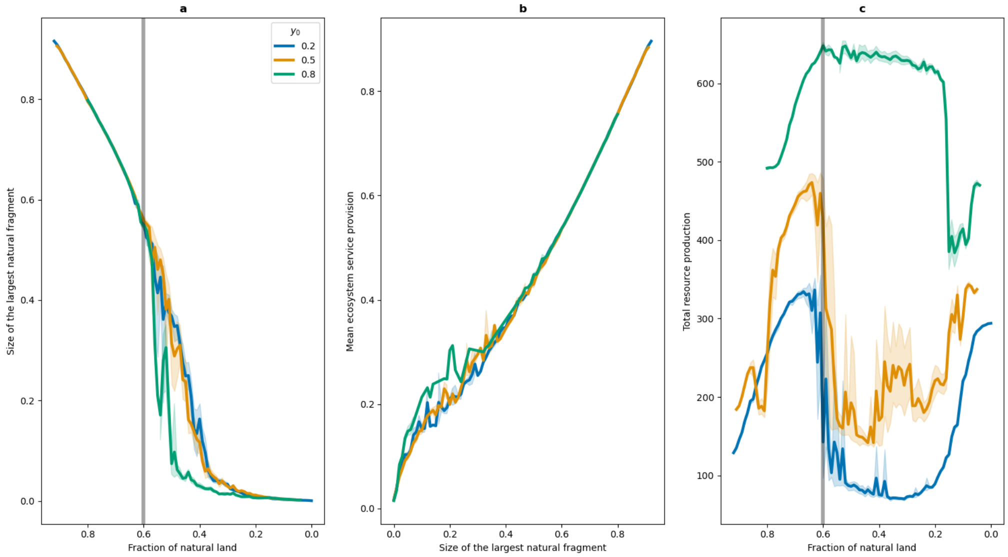


**Figure 4:** Landscape structure and consequences on ecosysetm service provision and agricultural production for different values of the ecosystem service saturation exponent. Solid lines are the mean of 5 replications and the shades represent the 95% confidence interval. a) Size of the largest natural fragment as a function of the fraction of natural land. The large decrease on the size of the largest fragment at the percolation transition is the same no matter the value of $z$. b) Baseline production does not have an effect on the relationship between mean ecosystem service and the size of the largest natural fragment. c) Increasing baseline production of low-intensity agricultural land reduces its dependence on ecosystem service provision and thus increases the resilience of agricultural yields against the fragmentation occuring after the percolation transition. In these simulations: $a = 0$, $w = 0$, $\sigma = 10$, $a_0 = 0.2$ and the rest of the parameters are set to the value presented on the main text table.

provision $\rho_D \simeq 0.02$ meaning that the average degradation time at null ecosystem service provision is of 100 years.

Finally, we assumed that fertility loss happens at the same timescales than degradation and chose to have an identical value for both parameters $\rho_D = \rho_L$. Large sensitivty of fertility loss with respect to ecosystem service provision shortens the average life time of agricultural cells and eventually agricultural patches do not persist enough time for them to provide resources to Humans thus causing population decrease or complete stagnation of population density growth (Figure 8). In the limit we placed ourselves, the average fertility loss is not a limitation to population growth in landscapes where natural land is very abundant.

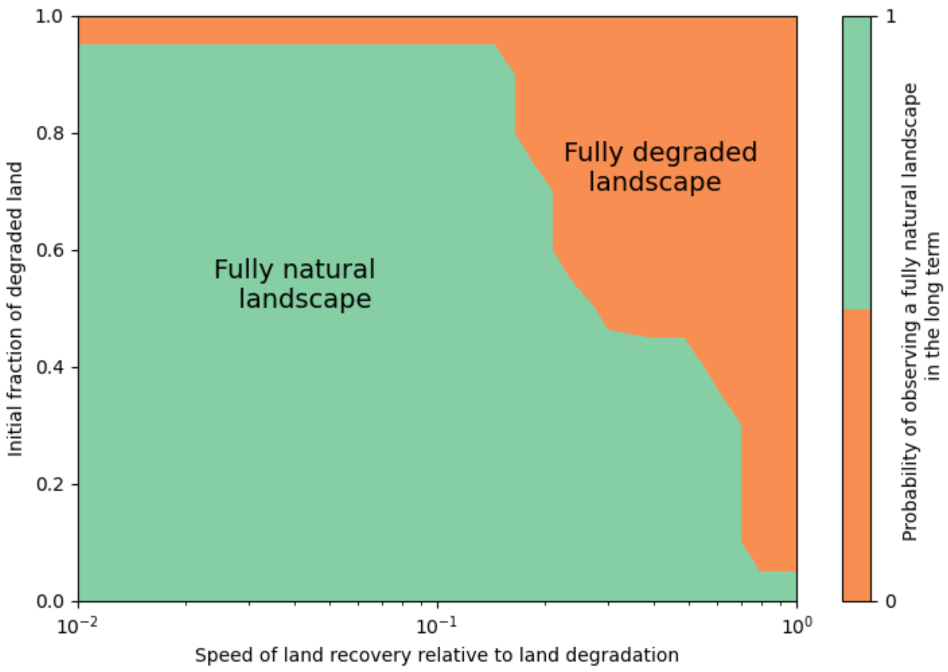


**Figure 5:** Final landscape state in the absence of Humans as a function of the initial fraction of degraded land and the relative speed of land recovery compared to land degradation respective to ecosysetm service provision. Choosing the recovery time at maximum ecosystem service provision to be 10 times smaller than degradation time at null ecosystem service provision place the landscape in a scenario where even at a 90% of degradation is fully recovered in the loing-term.

### Appendix 3: Simulation algorithm

Our model has an hybrid structure where the deterministic dynamics of Human population density are coupled to the stochastic dynamics of a landscape modelled as a cellular automata (i.e. with discrete spatialisation). Formulating landscape dynamics via the transition propensities between different types of land cover allow us to use the Stochastic Simulation Algorithm developped by (Gillespie, 1977) that yields exact realizations of the Stochastic Process (i.e. non-approximated solutions of the stochastic process master equation) in continuous time. In a nutshell, Gillespie's algorithm exploits the knowledge of the distribution probabilities of each transition to choose via drawing a random number the next time at which a transition occurs and the type of transition that occurs. This results in the simulation of every transition ocurring as time in the simulation is always advanced to the time at which a transition occurs. This simulation scheme is in conflict with the numerical solving in discrete time of the population dynamics ODE. We developped an hybrid

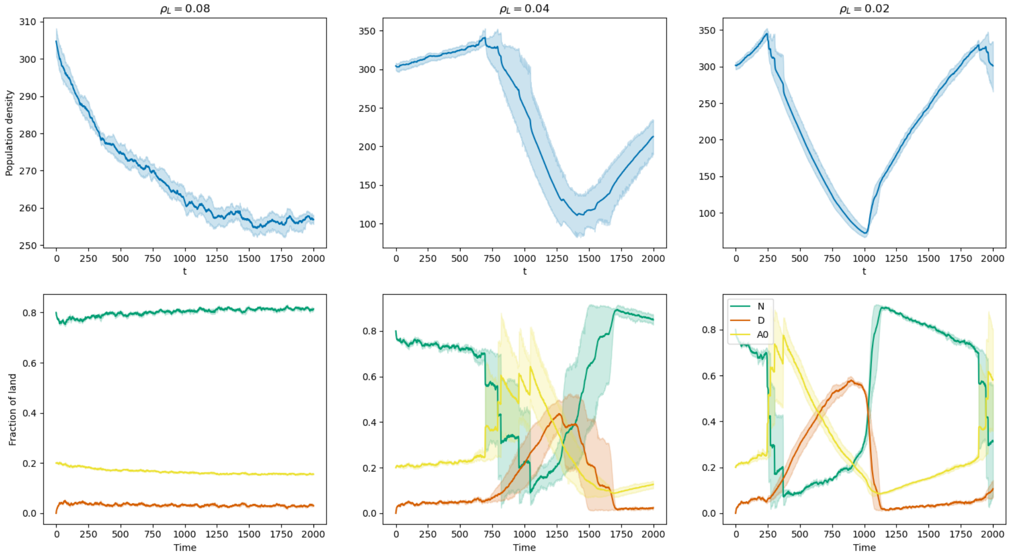


**Figure 6:** Social-ecological dynamics for different values of the sensitivity of fertility loss propensity per unit of ecosystem service. Solid lines are the mean of 5 replications and the shades represent the 95% confidence interval. Large sensitivity causes population decrease in a landscape with abundant natural land as the lifespan of agricultural cells is to short with respect to demographic processes making population density growth impossible. As the sensitivity increases, the percolation transition occurs before in time and with higher amplitude. The frequency of the collapse-recovery cycles decreases as the baseline production $y_0$ increases. In these simulations: $a = 0$, $w = 0$, $\sigma = 10$, $a_0 = 0.2$ and the rest of the parameters are set to the value presented on the main text table.

Stochastic Simulation Algorithm to deal with this conflict. In the hybrid version, the population dynamics ODE is solved at fixed time-steps, but all the transitions occurring between these timesteps are simulated. This is done by comparing each time the Gillespie timestep (which is the time until the next transition) and the demography timestep to assess what occurs first. When the Gillespie time-step is larger, population density is adjusted and in the opposite case the transition is simulated. Pseudo-code for the hybrid simulation algorithm is provided below in Algorithm 1.

### Appendix 4: Sensitivity Anlysis

We performed the model's exploration with OpenMole platform (https://github.com/openmole/openmole). We carried on a Morris Sensitivity Analysis to determine the relative importance of different parameters

**Algorithm 1** Hybrid Gillespie algorithm

Initialize population size, landscape and parameters;

Initialize simulation time $T$ and time-step for population ODE $dtp$;

$dt \leftarrow dtp$

**while** $t < T$ **do**

  Calculate the probability per unit time of each transition to occur on each patch;

  Generate a random number to calculate the time until next transition $d\tau$;

  **if** $d\tau < dt$ **then**

    Generate a random number to choose which transition will occur;

    Change the landscape according to the chosen transition;

    $t \leftarrow t + d\tau$;

    $dt \leftarrow dt - d\tau$;

  **if** $d\tau \geq dt$ **then**

    Solve the population size ODE

    $t \leftarrow t + dt$

    $dt \leftarrow dt_p$

on the model's outputs as well as the model robustness to changes in some parameters. The sensitivity analysis showed that the parameters that have the greatest influence on the model's outputs are the degree of agricultural clustering $\omega$, the baseline yield of low-intensity agriculture $y_0$ and the fertility loss $\rho_L$ and land recovery $\rho_R$ sensitivities to ecosystem service provision. The importance of the influence of $\rho_R$ and $\rho_L$ was however trivial. On the one hand allowing very fast (very slow) land recovery drastically diminishes (incereases) the likelihood of observing collapses. On the other hand and as discussed in Appendix 2, very fast fertility loss can prevent extremely unsustainable land-use management strategies from causing a collapes since the rapid loss of agricultural land prevents population growth. On the contrary, if fertility loss is very slow, agricultural land will accumulate and population grow enormously dramatically increasing the likelihood of a social-ecological collapse.

The large importance of $y_0$ on the model outputs' was also expected since increasing $y_0$ has the role of both increasing low-intense agricultural production but also decoupling it from ecosystem service provision making it inmune to habitat loss and fragmentation. The sensitivity analysis shows that increasing $y_0$ has positive effects both on human population size and the amount of natural land, as well as on containing degraded land. In terms of real-world interpretation this highlights the enormous importance that sustainable intensification, i.e. achieving high-yields with wildlife-friendly low-impact agriculture, will have on future sustainability (Bommarco et al., 2013). The importance of agricultural clustering in the model's outputs was less trivial but is completely understood following the model exploration. Our work highlights the importance of maintaining large connectrd natural fragments in agricultural landscapes to ensure ecosystem service provisioning by avoiding an habitat percolation transition and following massive fragmentation. Increasing $\omega$ has the effect of concentrating habitat loss in smaller regions of the space allowing for the persistence of very large natural fragments in the landscape and contributing to sustainability.

The difference of the impact between these parameters and the rest is not however huge, and no single parameter completely dominates model behaviour (see Figure 7). As we showed in the main-text, Humans' sensitivity to resource deficit $\sigma$ and the preference for agricultual intensification $\alpha$ can explain the appearence or disappearence of reversible collapses and sustainable steady-states. When we exclude $\rho_R$ and $\rho_L$ from the analysis (as their influence is trivial), the importance of $\sigma$ and $\alpha$ becomes more apparent.

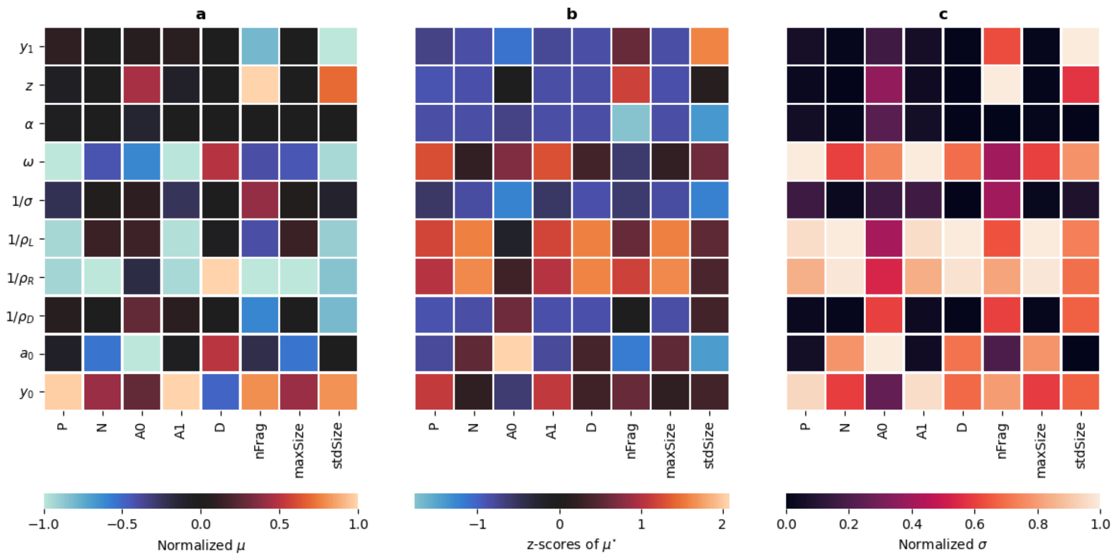


**Figure 7:** Sensitivty measures from the Morris sensitivity anlysis. Each row corresponds to a non-dimensional model parameter and each column to a measured output: nFrag is the number of natural fragments, maxSize the size of the largest fragment and stdSize the standard deviation of the fragment sizes. a) Normalized average of the elementary effects of each parameter on each ouptut. A positive value means that changes in the parameter value cause changes in the same direction for the outputs. A negative value means that the changes occur on the opposite direction. By construction this measure can have a value of zero if large effects on both directions are compensated. b) Z-score of the average of the absolute values of the elementary effects of changes of paramters on the model's outputs. This measure reflects the overall intensity of the effect of each parameter on the outputs of the model. Positive (negative) values mean that the parameter has an effect larger (smaller) than the mean of the effects. c) Normalized standard deviation of the elementary effects of changes in a model parameter on the outputs. This measure captures the level of non-linearity of each parameter on the outputs. A value of zero means that changes in a paramter have a linear effect on the output. The larger the value the more non-linear are the effects.

### Appendix 5 : Complement to Figure 1 of the main text

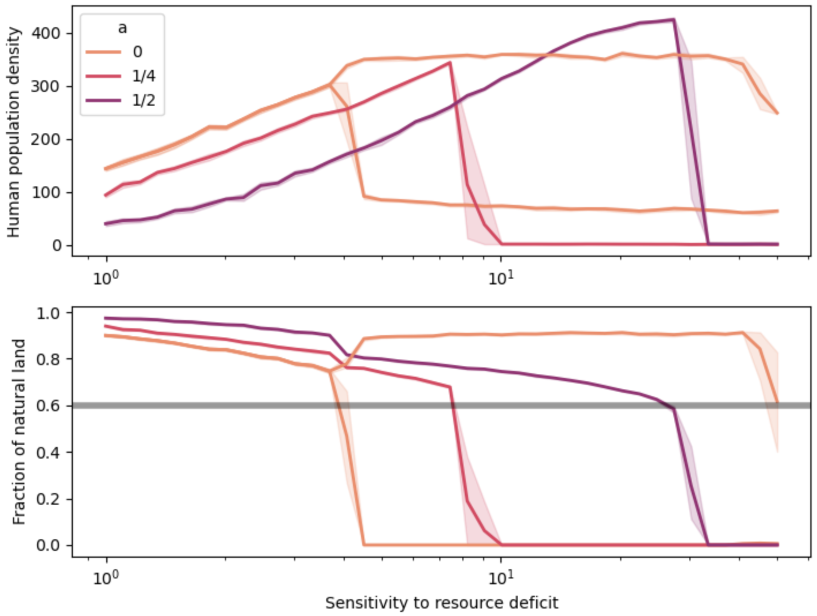


**Figure 8:** Bifurcation diagram for the human population density and the fraction of natural land as a function of control parameter $\sigma$ the sensitivity to resource deficit and for different values of the preference for agricultural intensification $a$. Solid lines are the mean of 5 replications and the shades represent the 95% confidence interval. The grey line in the bottom plot represents the percolation threshold for the natural fraction of land. Increasing the preference for intensification increases the resilience of the system with respect to the sensitivty to resource deficit, but however makes the collapes irreversible. For $a = 0$ the high and low lines after the bifurcation ($\sigma \simeq 4$ represent the cycles highest and lowest values respectively. In these simulations: $a = 0$, $w = 0$, $a_0 = 0.1$ and the rest of the parameters are set to the value presented on the main text table.
